# Bivariate genome-wide association analysis strengthens the role of bitter receptor clusters on chromosomes 7 and 12 in human bitter taste

**DOI:** 10.1101/296269

**Authors:** Liang-Dar Hwang, Puya Gharahkhani, Paul A. S. Breslin, Scott D. Gordon, Gu Zhu, Nicholas G. Martin, Nicholas G. Martin, Danielle R. Reed, Margaret J. Wright

## Abstract

Human perception of bitter substances is partially genetically determined. Previously we discovered a single nucleotide polymorphism (SNP) within the bitter taste receptor gene *TAS2R19* on chromosome 12 that accounts for 5.8% of the variance in the perceived intensity rating of quinine, and we strengthened the classic association between *TAS2R38* genotype and the bitterness of propylthiouracil (PROP). Here we performed a genome-wide association study (GWAS) using a 40% larger sample (n = 1999) together with a bivariate approach to detect previously unidentified common variants with small effects on bitter perception. We identified two signals, both with small effects (< 2%), within the bitter taste receptor clusters on chromosomes 7 and 12, which influence the perceived bitterness of denatonium benzoate and sucrose octaacetate respectively. We also provided the first independent replication for an association of caffeine bitterness on chromosome 12. Furthermore, we provided evidence for pleiotropic effects on quinine, caffeine, sucrose octaacetate and denatonium benzoate for the three SNPs on chromosome 12 and the functional importance of the SNPs for denatonium benzoate bitterness. These findings provide new insights into the genetic architecture of bitter taste and offer a useful starting point for determining the biological pathways linking perception of bitter substances.

## Introduction

Bitterness is a taste sensation that arises when particular chemicals come into contact with receptors in specialized cells on the human tongue^1-3^. But not everyone perceives the same bitterness for a given stimulus; this individual variation is partially genetically determined and can affect food perception, preferences and intake^4-6^. Genetic effects for bitter taste perception, which are estimated by twin studies, range from 36 to 73%^7-9^, with most of the known variation arising from inborn variation in the bitter receptor gene family (T2R)^10-12^. These bitter receptors are in tissues beyond the tongue and oral cavity, including the airways, gut, thyroid, and brain^13^ where they may function as toxin detectors or early-stage sentinel systems. Bitter taste responses may reflect how well the receptors detect ligands in other tissues^14^. Historically, the ability to taste one well-studied bitter compound, phenylthiocarbamide (PTC), has been related to many diseases^15^; more recently and more specifically, variation in the PTC taste receptor is shown to be involved in the immune system^16^ and to predict surgical outcome for severe rhinosinusitis^17^. Thus, together with the better-known effects on food intake and nutrition, bitter taste perception is of increasing importance to the medical field.

Given the rising importance of taste genetics, studies have focused on determining the underlying genetic variation that leads to individual differences in bitter perception. Our earlier genome-wide association study^12^ (GWAS), which included 1457 adolescents from 626 twin families, revealed a single nucleotide polymorphism (SNP) within the bitter taste receptor gene *TAS2R19*, accounting for 5.8% of the variance in the perception of quinine, and replicated the classic association between the bitter taste receptor gene *TAS2R38* and the perception of propylthiouracil (PROP; a chemical relative of PTC). The study, however, could neither detect loci for the other compounds tested, such as caffeine and sucrose octaacetate (SOA), that are likely to be affected by a large number of small-effect alleles nor the previously proposed but yet to be identified second locus for thiourea-containing compounds like PTC and PROP^18^.

Drawing on studies of complex traits such as body mass index^19^ and schizophrenia^20^, here we increased the overall sample size by 40% and used multivariate association analysis^21^ to identify common genetic variants (minor allele frequency [MAF] ≥ 5%) with small effects. Multivariate GWAS has been used to detect SNP associations that did not reach genome-wide significance in univariate analyses, such as autism spectrum disorders^22^ and bone mineral density^23^. This method can detect not only pleiotropic genetic variants but also variants associated with only one of the correlated phenotypes^24^. As shown by Stephens^24^, bivariate analysis increases power when there is greater separation of genotype groups (0, 1 or 2 copies of the minor allele) in two-versus one-dimensional space. In Figures 1a and 1b, we provide two illustrations of when a joint analysis of two correlated traits can provide greater separation of genotypes associated with the primary trait (trait 1). The first example (a) shows the case where only one trait (trait 1 on the y-axis) is associated with the variant (non-pleiotropic), with bivariate analysis providing better separation of the genotype groups in 2-dimensional space compared with the y-axis alone. A similar boost in signal would be found in a conditional analysis, where the non-associated trait is included as a covariate, as this removes the non-associated part of the variance in the associated trait (i.e., covariance between two traits) and, therefore, enhances the association. The second example (b) shows that maximum separation can be achieved when both traits (trait 1 on the y-axis, trait 2 on the x-axis) are associated with the variant and the effect of the minor allele on the two is in opposite direction. In the case where a variant has the same effect on both correlated traits (Figure 1c), bivariate analysis provides minimum/no increase in power. The bivariate approach is especially well-justified for bitter taste traits because, with the exception of PROP, perception of these bitter substances are highly correlated (r_p_ = ~0.6)^25^ and their genetic variances largely overlap (r_g_ = ~0.7)^7,9^.

**Figure 1.**
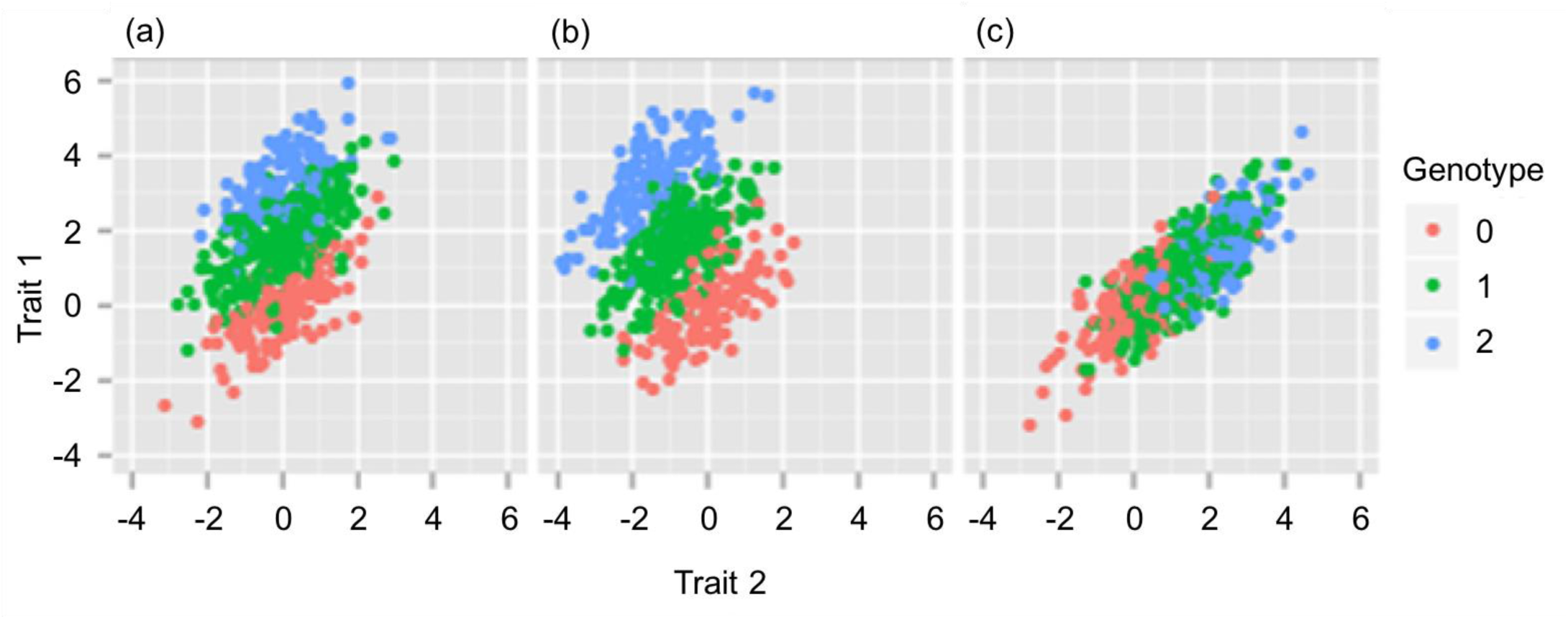
Illustration of three scenarios in a bivariate analysis. Each dot represents an individual, colored according to their genotype (0, 1 or 2 copies of the minor allele). In (a) trait 1 and 2 are correlated but the variant is only associated with trait 1. When considering traits 1 and 2 jointly in testing for association, there is greater separation of the genotype groups for trait 1 in the two-dimensional space compared with the y-axis alone. For example, the blue and green dots would largely overlap in the one-dimensional space along the y-axis. In (b) the minor allele has opposite effects on traits 1 and 2 - increasing trait 1 and decreasing trait 2. The three genotype groups are better separated in the two-dimensional space than for either trait individually. In (c) the minor allele has a similar effect on traits 1 and 2 - increasing both traits. Separation of the three genotype groups in two-dimensional space is no greater than along the y-axis alone. The figures and text are adapted from Figure 1 in Stephens (2013)^24^

Here we aimed to identify common genetic variants with small effects (i.e. 1-5%) on the perception of bitterness, building on our previous GWAS^12^, which was too under-powered to detect common genetic variants with small effects. We performed univariate GWAS for the perceived intensity of 5 bitter substances (PROP, quinine, caffeine, SOA, and denatonium benzoate [DB]) using our expanded sample, including 1999 individuals from 929 twin families. As these phenotypes were collected from the same individuals, to boost power we ran a series bivariate GWAS (6 in total) for the correlated phenotypes of quinine, caffeine, SOA and DB^9^. We looked for evidence of pleiotropy for each identified variant. When there was little evidence for pleiotropy, we tested the SNP association to the primary trait conditional on the second. For variants in linkage disequilibrium, we used bidirectional conditional analysis (i.e., including the genotype of one SNP as a covariate at a time to test the association with the other SNP) and plotted the SNP associations for one trait against the other. Finally, to help interpret the genotype-phenotype associations, we examined the potential function of the identified SNPs with bioinformatics tools.

## Results

We confirmed two previously identified associations with large effects on PROP and quinine, provided the first independent replication of an association for caffeine, and revealed two new associations with small effects (< 2%) on SOA and DB (Table 1). In addition, we found evidence for pleiotropic effects on quinine, caffeine, SOA and DB.

**Table 1.** Genetic variants associated with human bitter taste perception.

| Trait 1 | SNP | Chr:Position | A1/A2 | MAF | $\beta$ | SE | $r^2$ | P | Trait 2 | | | |
| --- | --- | --- | --- | --- | --- | --- | --- | --- | --- | --- | --- | --- |
|  |  |  |  |  |  |  |  |  | Quinine | Caffeine | SOA | DB |
|  |  |  |  |  |  |  |  |  | P_bivariate |  |  |  |
| Quinine | rs10772420 | 12:11174276 | G/A | 0.469 | -0.337 | 0.034 | 5.67% | **7.8e-23\*** | - | **4.8e-65\*** | **1.8e-24\*** | **6.4e-26\*** |
| Caffeine | <u>rs2597979<sup>†</sup></u> | 12:11189966 | G/C | 0.163 | 0.264 | 0.048 | 1.91% | **4.2e-8** | **8.4e-24\*** | - | **2.8e-10\*** | **4.5e-11\*** |
| SOA | <u>rs67487380</u> | 12:11194384 | A/G | 0.275 | -0.202 | 0.040 | 1.63% | 3.8e-7 | **5.4e-13\*** | **4.5e-8** | - | 2.4e-6 |
| DB | <u>rs10261515</u> | 7:141398707 | G/A | 0.491 | -0.136 | 0.037 | 0.93% | 2.5e-4 | **3.1e-8** | 4.0e-6 | 5.6e-4 | - |
| PROP solution | rs10246939 | 7:141672604 | C/T | 0.443 | 0.968 | 0.028 | 46.20% | **2.8e-199\*** |  |  |  |  |
| PROP paper | rs10246939 | 7:141672604 | C/T | 0.441 | 0.534 | 0.032 | 14.08%<sup>a</sup> | **5.4e-59\*** |  |  |  |  |
| PROP paper | <u>rs6761655<sup>‡</sup></u> | 2:218218646 | G/A | 0.186 | -0.246 | 0.044 | 1.83% | **2.7e-8** |  |  |  |  |

We report the top SNP from the peak association. SNPs that were not identified in our previous GWAS are underlined. Allele frequency and effect sizes are reported with reference to allele A1. Base-pair position is based on GRCh37; A1/A2, minor/major allele; MAF, minor allele frequency; $\beta$ , the effect size; SE, standard error of the $\beta$ ; $r^2$ , percent variance of the trait accounted for by the SNP; P, P-value from the univariate association analysis of trait 1; P\_bivariate, P-value from the bivariate association analysis of traits 1 and 2; SOA, sucrose octaacetate; DB, denatonium benzoate; **bold**, $P < \text{genome-wide significance threshold of } 5.0\text{e-}8$ ; \*, $P < \text{corrected significance threshold of } 1.0\text{e-}8$ ; <sup>†</sup>, an independent replication; <sup>‡</sup>, no evidence of replication. See Supplementary Tables 1 – 7 for the full list of SNPs.
<sup>a</sup> rs10246939 accounted only a third of the variance in PROP paper compared to PROP solution. This was partly due to the lower heritability of PROP paper ( $h^2 = 0.40$ ) compared to PROP solution ( $h^2 = 0.71$ , Supplementary Table S8).

### Confirmation of the locus on chromosome 12 influencing quinine and pleiotropic effects

The peak association for quinine was a missense variant within the bitter taste receptor gene *TAS2R19* on chromosome 12 (rs10772420, Figure 2a). As expected, with the increase in sample size the association was stronger (P = 7.8e-23) than that found in our initial GWAS (P = 1.8-e15)^12^, and the peak SNP explained almost the same amount of variance (5.67%). In the bivariate analysis, which included caffeine, there was a further boost in signal (P = 4.8e-65, Table 1). This was due to the nominal association of caffeine with rs10772420 (P = 2.5e-3; Figure 2b) and the effect of the minor allele being in the opposite direction to quinine (i.e. decrease in caffeine versus increase in quinine perception), which provided greater separation of the rs10772420 genotypes in two-dimensional space (as illustrated in Figure 1b). A much smaller increase in the quinine signal was found in the bivariate analysis with SOA (P = 1.8e-24) and DB (P = 6.4e-26). Both compounds (SOA: P = 1.0e-4; DB: P = 2.8e-3) were nominally associated with rs10772420 (Figures 2c and d), but the effect of the minor allele was in the same direction as that for quinine, resulting in little/no further separation of the genotypes in two-dimensional space (as illustrated in Figure 1c). Notably the size and direction of the effect of rs10772420 on the four bitter substances varied (Supplementary Figure 1; Supplementary Table 9): the strongest effect was on quinine (β= -0.337; 5.67% of the variance or 12.32% of the genetic variance), with a smaller fraction of the variance being explained for caffeine (β = 0.107; 0.57/1.24% of the total/genetic variance), SOA (β = -0.137; 0.94/2.04% of the total/genetic variance) and DB (β = -0.106; 0.56/1.22% of the total/genetic variance). In Figure 3 we show that variants with the largest effect on quinine - a cluster of 263 SNPs-were also associated with SOA, caffeine and DB, and that this cluster was separate to the top SNPs for SOA (a cluster of 167 SNPs) and caffeine (a cluster of 116 SNPs).

**Figure 2.**
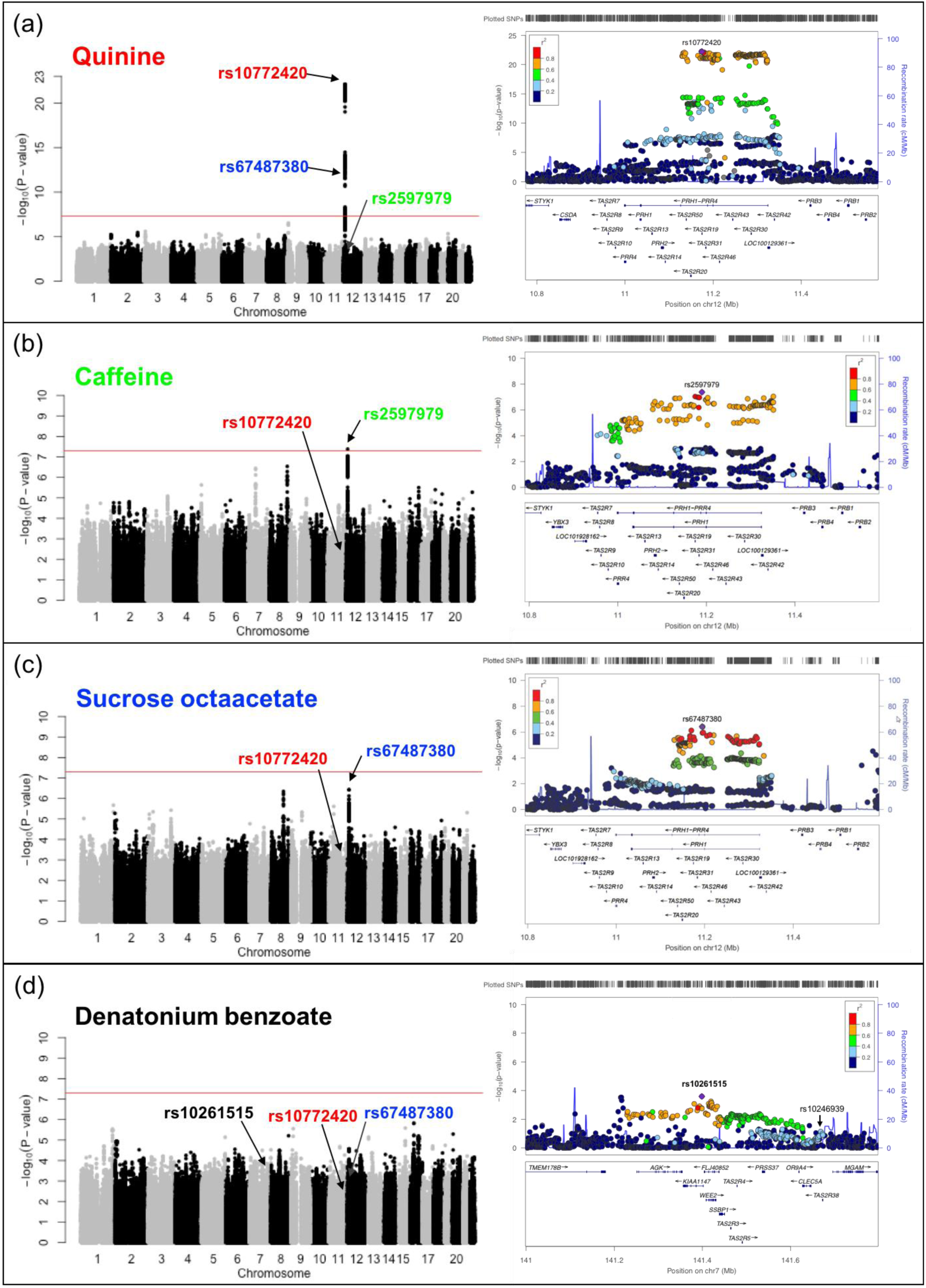
Common variants associated with the perception of bitter taste: (a) quinine, (b) caffeine, (c) sucrose octaacetate, and (d) denatonium benzoate (n = 1757) The left half of the figure shows the Manhattan plots, displaying the association P-value for each SNP in the genome (displayed as -log_10_ of the P-value). The red line indicates the genome-wide significance threshold of P = 5.0e-8. rs10772420 (labelled in red), rs2597979 (labelled in green), and rs67487380 (labelled in blue) are the most significant SNP within a putative or associated locus for quinine, caffeine, and sucrose octaacetate, respectively. rs10261515 is labelled in (d) because it reaches genome-wide significance in the bivariate analysis (Table 1 and Figure 4). The right half of the figure shows regional plots ±400kb for the top SNPs on chromosomes 12 (a, b and c) and 7 (d) with gene model below. Plots are zoomed to highlight the genomic region that likely harbors the causal variant. The top SNP for PROP (rs10246939) is also labelled in the regional plot in (d).

**Figure 3.**
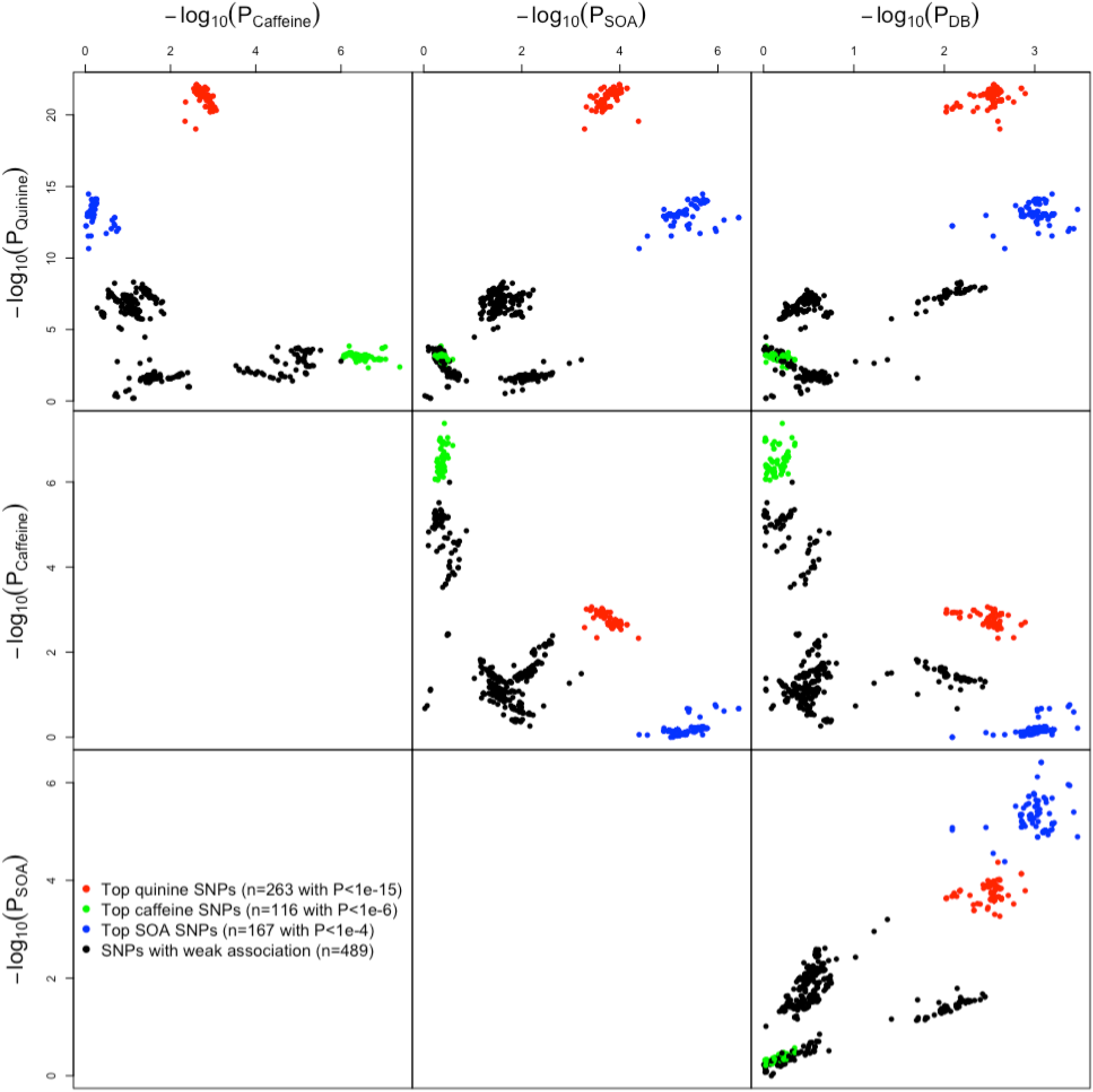
Top SNP associations on chromosome 12 for perceived intensity of quinine, sucrose octaacetate (SOA), caffeine and denatonium benzoate (DB) The red, blue and green clusters represent the top SNP associations with quinine, SOA and caffeine respectively. The top SNPs for these three bitter compounds are clustered separately from one another, even though the lead SNPs (rs10772420 for quinine; rs2597979 for caffeine; rs67487380 for SOA) of each cluster are correlated (r^2^_rs10772420-rs2597979_ = 0.24; r^2^_rs10772420-rs67487380_ = 0.43; r^2^_rs2597979-rs67487380_ = 0.08). The top SNPs for DB in this genomic region overlap with the tops SNPs for SOA, but the strengths of the associations with DB are weaker. In addition, there is evidence of pleiotropy. The red cluster is strongly associated with quinine, and more weakly associated with caffeine, SOA and DB; the blue cluster is associated with quinine, SOA and DB; the green cluster is associated with quinine and caffeine. A total of 1035 SNPs on chromosome 12 between 10950000 and 113550000 base pairs are plotted here.

### Independent replication of a SNP association on chromosome 12 for caffeine

For caffeine perception, we identified a peak association on chromosome 12 (rs2597979, P = 4.2e-8; Figure 2b), which accounted for a maximum trait variance of 1.91%. This SNP was in high linkage disequilibrium with that identified in a previous GWAS for caffeine detection threshold^11^ (r^2^ = 0.84 with rs2708377), and therefore we provide the first independent replication for this association. Further support was provided by our bivariate caffeine-quinine analysis (P = 8.4e-24). The enhancement in signal due to quinine also being associated with rs2597979 (P = 4.3e-3), with the effect in the opposite direction to caffeine (Supplementary Figure 1). Since the lead SNPs for caffeine (rs2597979) and quinine (rs10772420) were weakly correlated (r^2^ = 0.24), we tested whether the associations could be driven by the same SNP using conditional analysis, where each of the genotypes are included as a covariate. The caffeine-rs2597979 association remained (P = 4.4e-6; Table 2) after conditioning on the lead SNP for quinine, whereas the caffeine-rs10772420 association disappeared (P = 0.38) after conditioning on rs2597979, indicating that the caffeine-rs2597979 association was not driven by rs10772420. For quinine, the results of the conditional analysis were less clear. While the quinine-rs10772420 association remained highly significant after conditioning on the lead SNP for caffeine (P = 3.0e-19), a weak quinine-rs2597979 association remained after conditioning on rs10772420 (P = 0.044). Figure 3 shows that the top caffeine SNPs are weakly associated with quinine and largely independent from the top quinine SNPs.

**Table 2.** Conditional analyses of correlated SNPs on chromosome 12 associated with the perception of quinine, caffeine and sucrose octaacetate (SOA).

| Trait | SNP | Association (P-value) | Association conditional on correlated SNP (P-value) |  |  |
| --- | --- | --- | --- | --- | --- |
|  |  |  | rs10772420 | rs2597979 | rs67487380 |
| Quinine | **rs10772420** | 7.8e-23 | - | 3.0e-19 | 1.5e-10 |
|  | rs2597979 | 4.3e-3 | 4.4e-2 | - | - |
|  | rs67487380 | 1.5e-13 | 0.12 | - | - |
| Caffeine | rs10772420 | 2.5e-3 | - | 0.38 | - |
|  | **rs2597979** | 4.2e-8 | 4.4e-6 | - | 9.7e-8 |
|  | rs67487380 | 0.11 | - | 0.47 | - |
| SOA | rs10772420 | 1.0e-4 | - | - | 0.47 |
|  | rs2597979 | 0.38 | - | - | 0.63 |
|  | **rs67487380** | 3.8e-7 | 7.6e-4 | 5.3e-7 | - |

In contrast to quinine, we found little evidence for an association of either SOA or DB with rs2597979 (SOA: P = 0.38; DB: P = 0.62). The small boost in the caffeine-rs2597979 association found in the bivariate analysis (caffeine and SOA: P = 2.8e-10; caffeine and DB: P = 4.5e-11) was likely due to the correlation between the traits, which was supported by the boost in the caffeine-rs2597979 association when the intensity ratings for SOA (P = 5.9e-11) or DB (P = 7.9e-12) were included as a covariate. Figure 3 shows that the caffeine-associated SNPs are largely independent from SOA/DB-associated SNPs in this genomic region of chromosome 12.

### Putative novel associations identified in bivariate analyses influencing SOA and DB

The strongest association for SOA was found on chromosome 12 (rs67487380, P = 3.8e-7; Figure 2c). This SNP was also associated with quinine (P = 1.5e-13; Table 2, Figure 2a) and DB (P = 8.5e-4), with the size and direction of the effect being similar to that for SOA (Supplementary Figure 1), so that the boost in signal found in the bivariate SOA-quinine analysis (P = 5.4e-13; Table 1) was likely due to quinine. Even so, we found that the SOA-rs67487380 association remained when we conditioned on the lead SNP for quinine (P = 7.6e-4, Table 2), which is moderately correlated with rs67487380 (r^2^ = 0.43), whereas the SOA-rs10772420 association was lost (P = 0.47) when rs67487380 was included as a covariate. Similarly, for quinine, the rs10772420 association remained after conditioning on the lead SOA SNP (P = 1.5e-10), but the quinine-rs67487380 association disappeared (P = 0.12, Table 2), after conditioning on the lead quinine SNP. These conditional analysis results indicated that each of lead SNPs for SOA and quinine represents the main signal for its corresponding taste. Figure 3 clearly shows that the top SNPs for SOA and quinine are clustered separately from each other, whereas the top SNPs for DB in the genomic region on chromosome 12 largely overlap with the top SNPs for SOA.

In contrast to quinine and DB, caffeine was not associated with the lead SOA SNP (P = 0.11; Table 2). A small boost in signal in the bivariate SOA-caffeine analysis (P = 4.5e-8) was largely due to the correlation between SOA and caffeine. Further, the SOA-rs67487380 association remained after conditioning on the intensity rating for caffeine (P = 1.0e-8), indicating that the covariance between SOA and caffeine was not due to this SNP. Figure 3 shows that the top SNPs for SOA and caffeine are largely separated and this is because their lead SNPs are only subtly correlated (r^2^ = 0.08 between rs67487380 and rs2597979).

For DB, while all SNP associations had a P-value > 1.0e-6 (Figure 2d), one association on chromosome 7 (P = 2.5e-4) was boosted in the bivariate DB-quinine analysis (rs10261515, P = 3.1e-8, Table 1, Figure 4). The bivariate signal was mainly driven by the SNP association with DB, as there was no evidence for an association between quinine and rs10261515 (P = 0.15), and the DB signal was boosted after conditioning on the intensity score for quinine (P-value changed from 2.5e-4 to 1.9e-8). There was no evidence that this DB-SNP was associated with caffeine (P = 0.81) or SOA (P = 0.15), and little evidence of a signal boost in either the DB-caffeine (P = 4.0e-6) or DB-SOA (P = 5.6e-4) bivariate analyses (Table 1).

**Figure 4.**
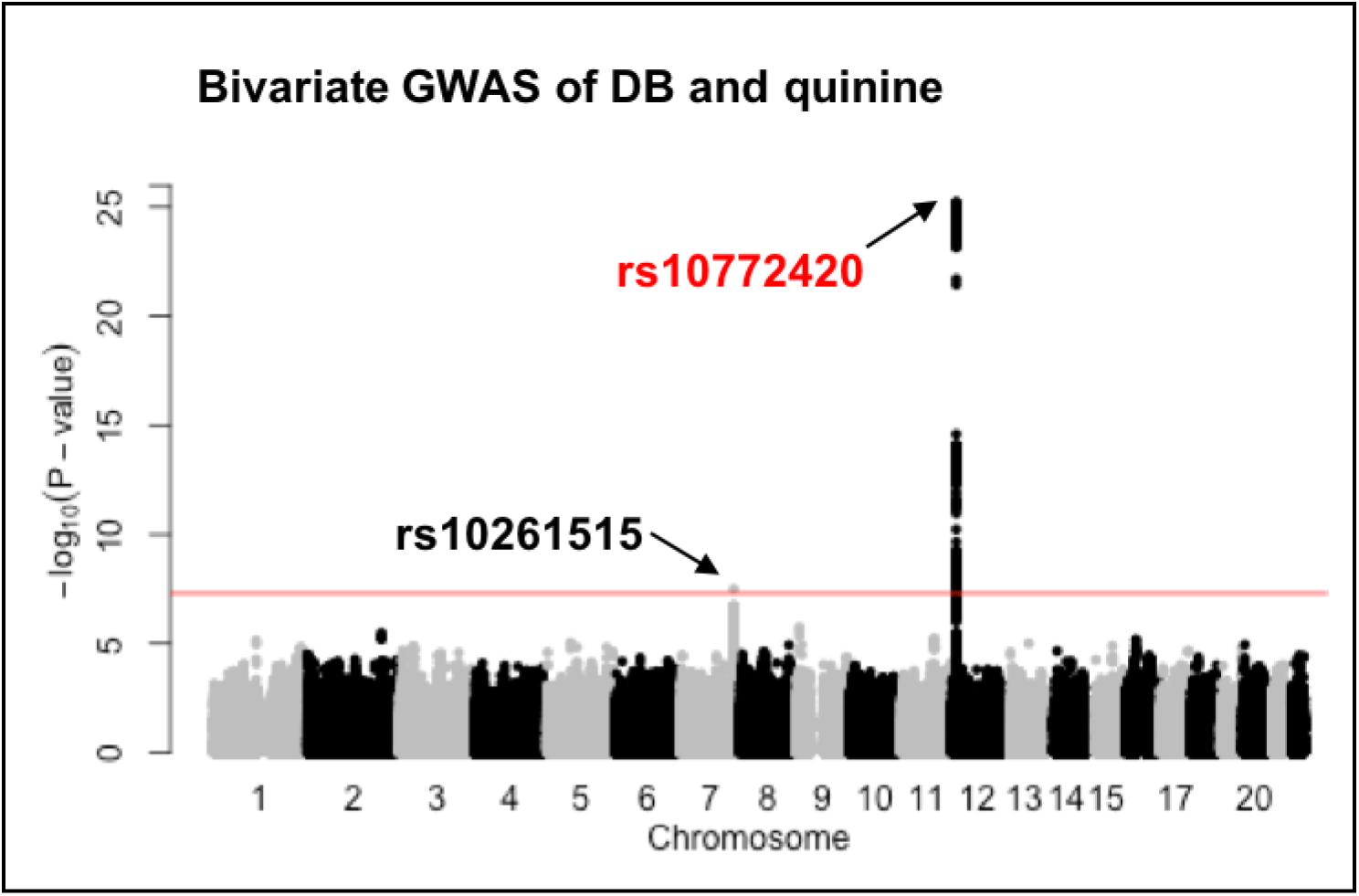
Manhattan plot showing a common variant (rs10261515) on chromosome 7 associated with the perception of denatonium benzoate (DB) based on the Bivariate GWAS of DB and quinine (n = 1757). The signal on chromosome 7 is driven by DB (P = 2.5e-4 in the univariate analysis) not quinine (P = 0.15). The signal on chromosome 12 is mainly due to the association of rs10772420 with quinine rather than DB as shown in Figures 2a and 2d. The red line indicates the genome-wide significance threshold of P = 5.0e-8.

The SNP rs10261515 is located within *KIAA1147* on chromosome 7, nearby three bitter taste receptor genes *TAS2R3*, *TAS2R4* and *TAS2R5* (Figure 2d), and is 274 kb upstream of the PROP-associated SNP rs10246939, with which it is weakly correlated (r^2^ = 0.23; Figure 2d). When we conditioned on the lead SNP for PROP, the DB-rs10261515 association remained (P = 9.0e-4), including after the additional adjustment for the quinine score (P = 1.7e-5).

### Confirmation of previously identified locus on chromosome 7 influencing PROP

The peak association for PROP was the well-known missense variant rs10246939 within the bitter taste receptor gene *TAS2R38* on chromosome 7 (Table 1, Supplementary Figure 2), confirming our previous findings^12^. For PROP paper, we identified a secondary locus within the *DIRC3* gene on chromosome 2 (rs6761655 and its completely correlated SNP rs6736242 [r^2^ = 1.0], P = 2.7e-8, Supplementary Figure 2b). This SNP accounted for a maximum trait variance of 1.83% in PROP paper and showed a weaker but nominally significant association with the perception of PROP solution (P = 7.4e-4). We note that this signal was present in our previous GWAS^12^ (Supplementary Figure 3), but was less obvious (i.e., not a solid peak – 4.4 M SNPs vs 2.3M SNPs in our earlier GWAS) and therefore was not reported. However, we found no evidence for this association with PROP perception in one previously reported GWAS of 225 Brazilians^10^, as well as two unpublished GWAS, one of ~500 individuals from the Silk Road population and one of ~2500 Italians (Supplementary Table 10). We further searched for this association in an earlier linkage study^26^, which prepared PROP paper in the exact same way as the present study, but the closest marker was ~500kb away from rs6761655 and it was not associated.

### Functional annotation of the identified SNPs

We performed functional analysis (i.e. the SNP effect on gene expression and DNA methylation) for five of the six SNPs in Table 1 using the bioinformatics tool Haploreg^27^. We did not include rs6761655 here due to lack of replication in the independent datasets. We also searched for bitter taste receptors that have been shown to respond to these bitter substances in human cell-based functional studies^28,29^. The key results are presented in Table 3 and a summary of the functional analysis can be found in Supplementary Table 11.

**Table 3.** Bioinformatics and cell-based functional studies of the genetic variants associated with bitter taste perception.

| Trait | Index SNP | GENCODE genes | Non-synonymous SNPs in LD ($r^2 \geq 0.8$) with index SNP | eQTL<sup>a</sup> | mQTL<sup>b</sup> | Cell-based functional analysis<sup>c</sup> |
| --- | --- | --- | --- | --- | --- | --- |
| Quinine | rs10772420 | TAS2R19 | rs10772420 in TAS2R19 | TAS2R10, TAS2R14, TAS2R19, TAS2R20, TAS2R31, TAS2R43, TAS2R50, TAS2R64P | TAS2R19 | T2R4, T2R7, T2R10, T2R14, T2R31, T2R39, T2R40, T2R43, T2R46 |
| Caffeine | rs2597979 | PRR4, TAS2R31 | rs10743938 in TAS2R31 | TAS2R14, TAS2R15, TAS2R20, TAS2R31, TAS2R43, TAS2R45, TAS2R64P | TAS2R19, TAS2R50 | T2R7, T2R10, T2R14, T2R43, T2R46 |
| SOA | rs67487380 | PRR4 |  | TAS2R10, TAS2R12, TAS2R14, TAS2R15, TAS2R19, TAS2R20, TAS2R31, TAS2R43, TAS2R46, TAS2R64P |  | T2R46 |
| DB | rs10261515 | KIAA1147 |  | TAS2R4, TAS2R5 |  | T2R4, T2R8, T2R10, T2R13, T2R30, T2R39, T2R43, T2R46 |
| PROP | rs10246939 | MGAM, TAS2R38 | rs713598, rs1726866 and rs10246939 in TAS2R38 | TAS2R5, TAS2R38 |  | T2R4, T2R38 |

<sup>a</sup> The genotype of the index SNP and/or its correlated SNPs ( $r^2 \geq 0.8$ ) is associated with the expression of these bitter taste receptor genes.
<sup>b</sup> The genotype of the index SNP and/or its correlated SNPs ( $r^2 \geq 0.8$ ) is associated with the methylation of DNA fragments within these bitter taste receptor genes.
<sup>c</sup> Bitter taste receptors shown to respond to bitter substances in cell-based functional analysis using human embryonic kidney cells<sup>28, 29</sup>.

The SNPs for quinine (rs10772420) and PROP (rs10246939) are missense variants within *TAS2R19* and *TAS2R38* respectively. In addition, the caffeine-associated SNP (rs2597979) is highly correlated with a missense variant rs10743938 (r^2^ = 0.92) within *TAS2R31*. This SNP has two possible allele changes of T>A and T>G, leading to residue changes of Leu162Met and Leu162Val respectively. In the present study, only rs10743938:T>A passed quality control and its association with caffeine had a P-value of 1.1e-7 (Supplementary Table 2).

Further, the SNPs for quinine, caffeine and SOA are common expression quantitative loci (eQTL) for five bitter taste receptor genes (*TAS2R14*, *TAS2R20*, *TAS2R31*, *TAS2R43*, *TAS2R64P*) on chromosome 12, and the expression of other bitter taste receptors in the same region is regulated by one or two of these three SNPs, e.g., the expression of *TAS2R46* is only regulated by the SOA and quinine associated SNP rs67487380. The DB-associated SNP rs10261515 influences the expression of the bitter taste receptor genes, *TAS2R4* and *TAS2R5*, on chromosome 7. T2R4 is more likely to a receptor for DB because the allele (rs10261515 G allele) for weaker DB intensity rating is associated with a lower expression level of *TAS2R4* and the opposite for *TAS2R5*. In addition, DB can activate T2R4 but T2R5 in cell-based functional analysis. Results from the cell-based functional analysis do not necessarily agree with the results from the bioinformatics functional analysis. For example, the quinine-associated SNP rs10772420 is a missense variant within *TAS2R19* and it regulates both gene expression and DNA methylation of *TAS2R19*, but T2R19 does not respond to quinine. We note that neither of these bioinformatics and cell-based functional analyses were based on human taste tissues.

## Discussion

In this study of bivariate GWAS on human taste perception, we identify two putative novel associations, including rs67487380 on chromosome 12 for SOA-elicited bitter taste and rs10261515 on chromosome 7 for DB-elicited bitter taste. In addition, we provide the first independent replication of an association on chromosome 12 for caffeine bitterness (rs2597979) and confirm our previously reported associations for quinine bitterness (rs10772420 on chromosome 12) and PROP bitterness (rs10246939 on chromosome 7). All variants are located within the bitter taste receptor clusters on chromosomes 7 and 12, highlighting the importance of these two regions in the genetics of bitter taste. Further, we show evidence of pleiotropy for those variants on chromosome 12 and the functional importance of the DB-associated SNP.

This is the first GWAS study to identify a SNP (rs67487380 on chromosome 12) association with human perception of SOA. In mice, a major locus for SOA perception (*soa*) was reported in the early 1990s^30^. Interestingly, the mouse *soa* locus also affects the perception of other bitter substances, including quinine, DB, PROP, but not caffeine^31,32^. Here we provide evidence that rs67487380 is also associated with the perception of quinine and DB, but not caffeine or PROP (P > 0.05). SOA activates human T2R46 but no other T2Rs in heterologous expression assays^29^. It is possible that rs67487380 regulates the perception of SOA through its effect on mRNA expression because the G allele for weaker SOA intensity is also associated with a lower expression level of *TAS2R46*. Nevertheless, rs67487380 could still be a proxy for true causal variants.

The finding of the novel association between DB and the SNP rs10261515 suggests that there may be a second locus on chromosome 7 that affects human bitter taste perception (the first is the locus within *TAS2R38* for PROP). Heterologous expression studies using human embryotic kidney (HEK) cells transfected with *TAS2Rs* have shown that DB activates T2R4 but no other bitter taste receptors in this region (e.g. T2R3, T2R5 and T2R38)^28^. In addition, the human T2R4 is the ortholog of mouse T2R8, which also responds to DB^1^. Our functional annotation results provide further support for T2R4 as a DB bitter taste receptor, since the allele (rs10261515 G allele) for a lower perceived intensity of DB is associated with lower expression level of *TAS2R4* mRNA.

The SNP association for caffeine perception replicated a previous GWAS of 607 Brazilian adults^11^. In that study the lead SNP accounted for 8.9% of the variance in caffeine sensitivity, compared with our estimate of 1.9%. Similarly, the Brazilian study accounted for 23.2% of the variance of quinine taste with genetic mutations, which is four times the effect estimated here. This difference in effect sizes is likely due to two main factors. First, the taste scores in the Brazilian sample were corrected for overall-taste-sensitivity (an average score of the perception of sweet, umami, sour, salty and bitter compounds), which removed ~30% of the variance in the perception of caffeine and quinine. Without correction, rs10772420 accounted for 13.2% of the variance in quinine, and the caffeine association was not detected due to low power. Second, the Brazilian study used a detection threshold approach, which measures overall oral sensitivity, compared with our measure of bitter taste intensity. Regardless, both studies identified the same variants for caffeine, quinine as well as PROP, indicating that these are likely to be valid associations among human bitter taste perception and these T2R-rich regions of chromosomes 7 and 12.

The functional annotation of the caffeine-associated SNP showed that the highly correlated SNP (rs10743938) is a missense mutation that could affect the function of T2R31. This is the first evidence linking this bitter taste receptor to the perception of caffeine, while genetic variants in *TAS2R31* have been shown to affect the perception of quinine^33^ and acesulfame potassium^34^ (a non-nutritive sweetener with bitter aftertaste). Prior cell-based functional studies^28^ reported that caffeine does not activate T2R31 in heterologous expression assays; rather, it activates T2R7, -10, -14, -43, and -46, and that the summed expression level of these activated T2Rs increases with the perceived intensity of caffeine^35^. However, comparing results from bioinformatics and cell-based analyses can be limited by two major factors. Here, we report associations for the index (lead) SNP with the lowest P-value, but since this SNP is in a linkage disequilibrium block, the association could be driven by any variant within the block. Second, these cell-based functional assays^28^ were conducted in heterologous systems (i.e., HEK cells transfected with *TAS2Rs*), which may not always recapitulate human sensory experience faithfully^36^. We observed a similar difference for quinine, with the lead quinine-associated SNP rs10772420 constituting a missense mutation in *TAS2R19*. Yet T2R19 does not respond to quinine in functional expression assays^28^. Therefore, a better method to identify causal SNPs for the foreseeable future is to tightly integrate genetic-perceptual association results with those of taste receptor cell-based assays using human taste tissues, such as taste buds or cultured human taste cells^37^.

This study provides the first evidence for antagonistic genetic pleiotropy in bitter taste. The two SNPs rs10772420 and rs2597979 have opposite effects on the perceived intensity of quinine and caffeine (Supplementary Figure 1; Supplementary Table 9) and this largely enhances the strengths of their associations (P-value) in the bivariate analysis (Figure 1b). As bitter-tasting substances (e.g., caffeine) can have both beneficial and detrimental effects, the antagonistic pleiotropy may be an evolutionary consequence that avoids over and under consumption.

The top SNPs for quinine, caffeine, and SOA were correlated (r^2^ = 0.08 – 0.43) and each could have various effects on one another. These correlations are due to the linkage disequilibrium between polymorphisms within bitter taste receptor genes on chromosome 12, which results in common haplotypes for nearby genes and long-range haplotypes for more distant ones^38,39^. Previous studies have revealed a complex bitter substance – receptor relationship, with one bitter compound activating multiple T2Rs and one T2R responding to multiple bitter substances^28,29,40^. Taken together, it is likely that the perception of a bitter taste can be mediated by multiple T2Rs, and SNPs identified in the present study could represent haplotypes that regulate several T2Rs together. We have attempted to illustrate this in Figure 5 by taking the perception of quinine and caffeine as an example. The lead SNP for quinine (rs10772420) is correlated with several SNPs (the regional association plot in Figure 2a) that regulate the T2Rs for quinine (cell-based functional analysis results in Table 3). Also, the lead SNP for caffeine (rs2597979) is correlated with SNPs (Figure 2b) that regulate T2Rs for caffeine (Table 3). In addition, the common T2Rs for the two tastes are regulated by SNPs that are in linkage disequilibrium with the two lead SNPs. We note that the real regulatory network can be more complex than this, such that one T2R can be regulated by multiple SNPs. Whereas we used conditional analysis (Table 2) and plotted the SNP associations against the three tastes (Figure 3) to show that each of the lead SNPs represents the main signal in the linkage disequilibrium block, the clusters of nearby bitter receptors and many variants in high linkage disequilibrium create challenges in separating causal from non-causal variants.

**Figure 5.**
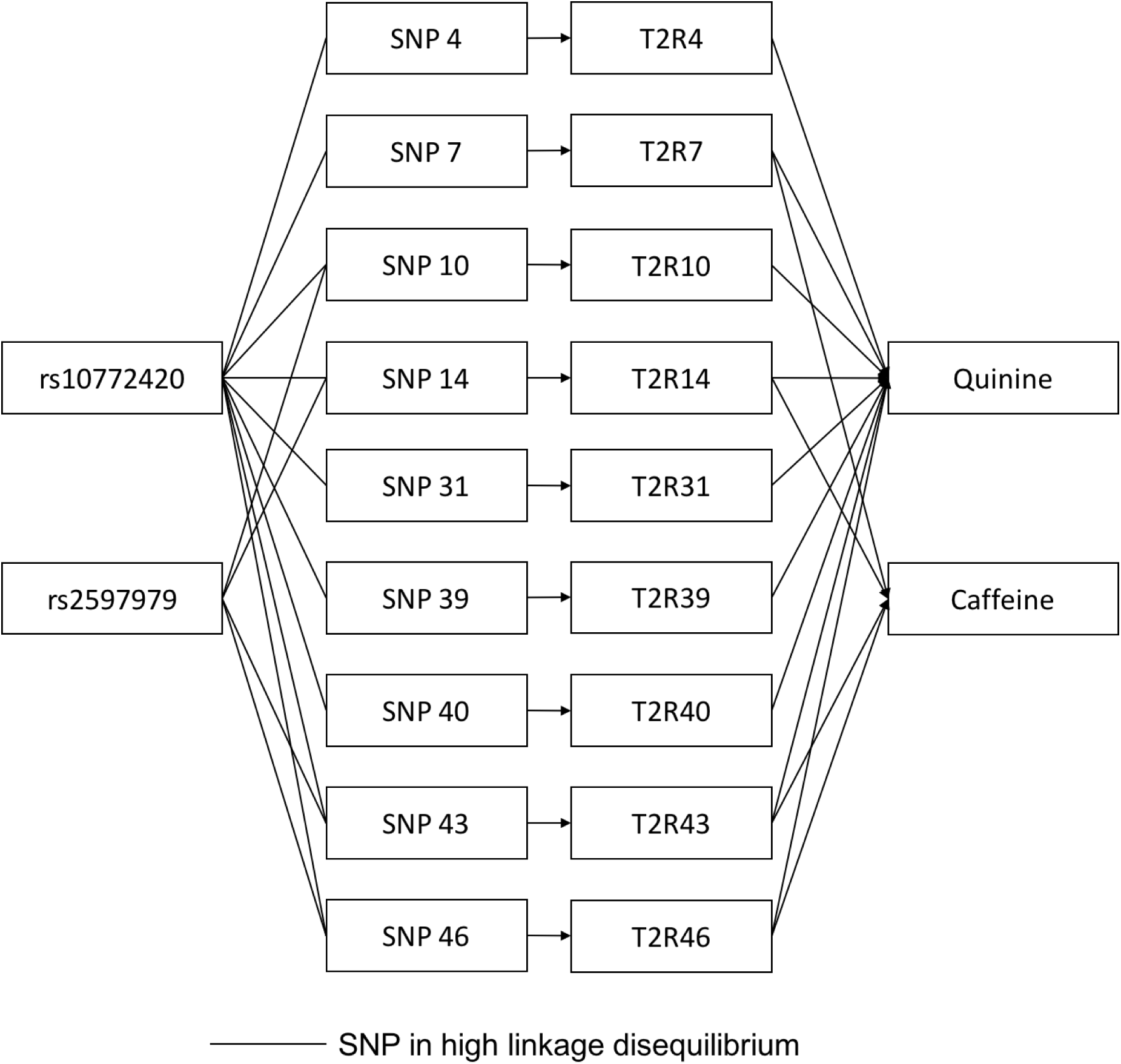
Potential model of the SNP regulation of human bitter taste perception. Quinine can be detected by bitter taste receptors T2R4, -7, -10, -14, -31, -39, -40, -43, and -46 on chromosome 12, and caffeine can be detected by the T2R7, -14, -43, and -46 (as summarized in Table 3), which overlap the T2Rs for quinine. Here we assume that each T2R is regulated by a major SNP with the corresponding number. rs10772420 is associated with the perception of quinine via its correlated SNPs; rs2597979 is associated with the perception of caffeine via its correlated SNPs.

Perceptual studies of bitter taste also have reported that individual differences in perceived bitterness from multiple compounds show positive correlations. Most relevant to the present work, past studies demonstrated a strong correlation of perceived bitter taste intensities among DB, SOA, and quinine^25^. This observation harkens to that reported in the present study for rs67487380 on chromosome 12. Furthermore, individual differences in bitterness from SOA, caffeine and quinine were also observed, suggesting a linkage between SOA receptor variants and caffeine receptor variants^25^. This too reflects associations observed in the present data set. Perhaps, a linkage disequilibrium block accounts both for the genetic architecture as well as the bitterness perception associations.

Prior work using pedigree segregation analysis has proposed that the perception of PTC (a structurally related chemical to PROP) is modulated by a 2-locus model^18^, but the location of a second locus has been unclear for nearly 30 years. Here we found no support for an association with *TAS2R1*, which was suggested by a prior family-based linkage study^26^, but identified a putative secondary locus within the *DIRC3* gene on chromosome 2, which accounted for an additional 1.83% of the variance (4.58% of the genetic variance) in the perception of PROP paper. While we found no evidence for replication using three independent datasets – from one published study (i.e. the Brazilian sample) and two unpublished (the Silk Road and the Italian samples), we note that there are considerable differences across studies (e.g., sample age (all other studies used adults), ethnicity, and delivery method (the Brazilian study used PROP solution)), which may have influenced our ability to replicate their findings. We did a further search for this association using available data from an early linkage study^26^, which used the same PROP paper, but were severely limited by the sparsity of markers, as none were close to the SNP identified in the present study. Ideally, we need to test for this association using the same methods and materials (i.e., the perceived intensity of saturated PROP paper measured from adolescents with European ancestry), but at this stage the signal does not appear to be sufficiently robust to be detected with alternative methods.

The strengths of the present study include the use of the largest sample to date, and the collection of multiple taste phenotypes from the same individuals, which increases the statistical power via bivariate association analysis. We show that the association signals (P-value) for quinine and caffeine (rs10772420 and rs2597979 respectively) were stronger in the bivariate compared with the univariate analysis, but the estimated effect size remains the same. The signal boosts in these already established associations serve as a proof of principle for using bivariate GWAS. We also show that, through the discovery of the association of DB, a signal can be enhanced when only one of two correlated traits is associated. This is useful for identifying non-pleiotropic SNPs for correlated phenotypes. We used multiple levels of analysis (conditional on genotype and phenotype) as well as cluster plots to disentangle the pleiotropic nature of these SNPs with bitter tastes and provide additional support for the signals identified in the bivariate analyses. We attempted to obtain data to replicate every novel association. However, we were unable to test the association for SOA and DB, due to no other datasets being available. Given the enhancement in the known signals for both quinine and caffeine in the bivariate analyses, together with the post-hoc bioinformatics analyses, as well as prior functional analyses, we believe the SOA and DB hits are unlikely to be false positives. Further, findings from multivariate GWAS of other phenotypes, e.g., levels of plasma lipids^41^, have been replicated in independent studies. The variants for SOA and DB account for less than 10% of the genetic variance (< 2% of trait variance) of their associated traits, suggesting that there are more variants with smaller effects. The remaining genetic variance could be partly due to rare variants because SNPs with an MAF smaller than 5% were excluded here and rare variants can have a large effect on complex traits^42^.

In conclusion, this study reveals the influence of multiple variants on bitter taste and demonstrates the benefits of multivariate analysis over the conventional univariate GWAS. Recent advancement in the methodology of multivariate GWAS (i.e. MTAG^43^) could make multivariate analysis easier to apply because it uses individual summary level results from different studies and does not require correlated phenotypes to be collected from the same sample. Whereas our previous twin analysis provided strong evidence of pleiotropy for the perception of several bitter compounds (except for PROP), there are numerous causal models that could underlie this shared genetic etiology. Identification of specific SNPs/genes involved offers a useful starting point for determining the biological pathways linking perception of bitter substances and for delineating of the mechanisms involved. Future studies integrating bioinformatics and functional analyses using human taste tissues will provide stronger evidence in identifying true causal variants, which could assist personalized nutrition and precision medicine.

## Materials and Methods

### Sample

Participants were 1999 adolescent and young adult Caucasian twins and their siblings from 929 families from the Brisbane Adolescent Twin Study^44^, also referred to as the Brisbane Longitudinal Twin Study (BLTS), with data collected between August 2002 and July 2014. This sample consisted of 275 MZ and 544 DZ twin pairs, including 155 pairs with one to two singleton siblings, and 184 unpaired individuals (mean age of 16.0 ± 2.8 years [medium 14.5 years, range 11-25years]; 1075 females, 924 males). It included all participants from our previous genome-wide association study^12^, plus a 40% increase in sample size.

### Taste Test

The taste test battery has been described in detail elsewhere^7^. Briefly, it included duplicated presentations of five bitter (6.0 × 10^-4^ M PROP, 2.0 × 10^-4^ M SOA, 1.81 × 10^-4^ M quinine, 0.05 M caffeine, and 4.99 × 10^-6^ M DB) solutions as well as a paper strip rinsed in a saturated PROP solution (0.059M). Participants were instructed to rate their perceived intensity for each solution and the PROP paper using a general Labelled Magnitude Scale (gLMS)^45^ with labels of no sensation (0 mm), barely detectable (2 mm), weak (7 mm), moderate (20 mm), strong (40 mm), very strong (61 mm), and strongest imaginable (114 mm). Mean intensity ratings from duplicate presentations for each stimulus were used in all analyses. A total of 1757 participants completed the full test battery (solutions and PROP paper) with a further 242 providing an intensity rating for the PROP paper only.

### Genotyping, Genetic Imputation and Quality Control

Genotyping was performed with the Illumina 610-Quad BeadChip (n = 1457 individuals) and the HumanCoreExome-12 v1.0 BeadChip (n = 542 individuals), with approximately 700k SNPs passing standard quality control filters, as outlined previously^12^. These SNPs were then phased using ShapeIT^46^ and imputed using Minimac3^47^ and the Haplotype Reference Consortium of Caucasian European ancestry (Release 1)^48^. Individuals who were > 6SD from the PC1/PC2 centroid were excluded, so our sample was of exclusively European ancestry. To ensure SNPs were imputed with high data quality, SNPs with a call rate < 90%, MAF < 0.05, imputation score < 0.3, and Hardy–Weinberg equilibrium score of P < 10^-6^ were excluded. Approximately 4.4M SNPs met these criteria and were used in the analyses.

### Genome-wide Association Analysis

Univariate and bivariate GWAS were conducted using GEMMA^21^, which fits a linear mixed model for each SNP and uses the genetic relatedness matrix to account for the family structure. Covariates included age, sex, a history of ear infection, all of which were shown to be associated with taste intensity ratings^9^, and the first five principal components calculated from the genotypes. Bivariate analysis essentially provides a complement to univariate analysis. It can enhance the strength of a SNP association, but the estimated effect on each of the two traits remains. For non-pleiotropic SNPs identified in bivariate analysis, we tested for their associations using conditional analysis of the associated trait conditional on the non-associated trait. When two identified SNPs were correlated, to test whether they were independent signals for the corresponding traits, we performed conditional analyses, by fitting each of the SNPs as an extra covariate. Prior to analyses, intensity ratings for each stimulus were square-root transformed to obtain a more normal distribution^9^ and then converted to Z-scores. A genome-wide significance threshold was defined as P < 5.0e-8. As four of the phenotypes were correlated (r_p_ between quinine, caffeine, SOA and DB = 0.58 – 0.64)^9^ the number of independent tests was estimated using a matrix spectral decomposition algorithm^49^ at 4.96 and accordingly a Bonferroni-corrected threshold was defined as P < 1.0e-8. The genomic inflation factor (λ) ranged between 0.99 and 1.02 (Supplementary Figures S4 and S5), which indicates that potential technical or population stratification artifacts had a negligible impact on the results. As all association analyses were performed under an additive model and all phenotypes were converted to Z-scores, variance explained by a SNP was calculated as 2 × *MAF* × (1 − *MAF*) × *β*^2^. Manhattan and Q-Q plots were created using the “fastman” package^50^ in R. Regional association plots were created using Locuszoom^51^.

### Functional annotation of the identified SNPs

To examine the potential role of the identified SNPs, we used Haploreg v4.1^27^ for functional annotation. Briefly, it annotates all index SNPs and their correlated SNPs (r^2^ was set to be > 0.8 for this study) by their associated chromatin states (e.g., conserved regions and DNAse hypersensitivity sites) from the Roadmap epigenomics project^52^ and Encode project^53^ and their effects on regulatory motifs. It also reports the effect of SNPs on gene expression in multiple tissues from eQTL (expression quantitative trait loci) studies, including results from the GTEx^54^ project portal. Use of functional annotation may provide more information about the putative role of a specific gene as well as developing mechanistic hypotheses of the impact of the SNP on phenotypes (e.g. variation in taste perception). More details are provided in Supplementary Table S3.

### Ethical Statement

The Queensland Institute of Medical Research Human Research Ethics Committee approved the study. Written informed consent was obtained from both the participants and their parents (the latter not required for those 18 years and over) before participation. All methods were performed in accordance with the relevant guidelines and regulations.

## Acknowledgements

We thank Kirsten J Mascioli, Christopher Tharp, Fujiko Duke, Deborah Lee and Corrine Mansfield from the Monell Chemical Senses Center for manufacturing the taste tests; and from the QIMR Berghofer, Marlene Grace, Ann Eldridge, Natalie Garden, and Kerrie McAloney for project co-ordination, data collection, and data entry, and David Smyth for computer support. We thank Robino Antonietta from Italian Ministry of Health for looking up SNPs in their unpublished GWAS of PROP. In particular, thanks go to twins and their families for their participation.

## Author Contributions

Conceived and designed the experiments: PASB NGM DRR MJW. Processed data: LH SDG. Analyzed data: LH PG GZ. Drafted the first version of the manuscript: LH. Provided critical feedback on the manuscript: PG GZ PASB NGM DRR MJW.

